# Sensitive untargeted metabolomics via 3-nitrophenylhydrazine derivatization maps gut microbiota-related metabolites across multiple mouse biological compartments

**DOI:** 10.64898/2026.10.27.762144

**Authors:** Qingqing Wu, Li Zhu, Yi Chi, jie Dai

## Abstract

The gut microbiome communicates with distal organs via small molecules, yet systematic tracking of gut microbiota-related metabolites across host compartments remains limited. Although 3-nitrophenylhydrazine (3-NPH) derivatization improves sensitivity for carboxyl-, carbonyl- and phosphate-containing metabolites, the resulting derivatives cannot be identified against conventional spectral libraries. Here, we developed a 3-NPH derivatization-based untargeted metabolomics approach coupled with a purpose-built theoretical spectral library and identified 301 gut microbiota-related metabolites across five compartments of young and adult mice of both sexes. Cross-compartment comparisons revealed compartment-specific compositions and fewer sex-associated differences in serum, liver and brain than in the gut. Spearman correlation analysis along the gut-brain and gut-liver axes identified coordinated metabolite pairs, including glycolysis intermediates and stereochemically diverse fatty-acid derivatives, supporting transmission of gut microbiota-related metabolites to circulation and distal organs. Our approach overcomes the long-standing identification bottleneck of derivatization-based untargeted metabolomics, providing a transferable platform for compartment-resolved mapping of gut microbiota-related metabolites and highlighting the gut-brain and gut-liver axes as routes for microbiota-host metabolite exchange.

## Introduction

The intestinal microbiota produces a vast and chemically diverse set of small molecules that influence host physiology far beyond the gut lumen^1^. Microbiota-derived metabolites, including short-chain fatty acids (SCFAs)^2^, amino acid derivatives, phenolic compounds, indoles and bile acids^3^, act both locally and distally, and are increasingly recognized as key mediators of the gut-brain and gut-liver axes^4,5^. Despite rapid progress^6^, a systematic, compartment-resolved view of where gut-derived metabolites are present, and how they are shared across the gut, circulation and peripheral organs, is still lacking.

A central obstacle is analytical. Many microbiota-relevant metabolites are polar, weakly ionizing compounds that are poorly detected by conventional reversed-phase liquid chromatography-mass spectrometry (LC-MS); chemical derivatization markedly improves their sensitivity^7^. However, derivatized products produce spectra that no longer match standard spectral libraries, making their identification difficult^8^. To address this, we established an untargeted metabolomics method based on 3-nitrophenylhydrazine (3-NPH) derivatization combined with a purpose-built theoretical spectral database, in which each candidate metabolite is computationally derivatized and a 3-NPH-specific diagnostic fragment ([C□H□N□O□]□, m/z 137.035) is enforced to lock identification.

In this study, we established and applied the derivatization-based untargeted metabolomics platform to quantify gut microbiota-related metabolites in five biological compartments (cecum contents, feces, serum, liver and brain) from young (2-3-week) and adult (8-10-week) mice of both sexes. We profiled the coverage and composition of the detected metabolome in each biological matrix, delineated age- and sex-associated differences, and used Spearman correlations on within-mouse, compartment-paired samples along the gut-brain and gut-liver axes to explore potential transmission routes from the gut to the systemic circulation and distal organs such as the brain and liver.

## Results

### Establishment of a 3-NPH derivatization-based untargeted metabolomics method

We established a 3-NPH derivatization-based untargeted metabolomics method. The analytical workflow of this method is illustrated in Fig. 1a. Briefly, every derivatizable metabolite is computationally processed and queried against a purpose-built theoretical spectral database. For each candidate bearing at least one carboxyl, carbonyl or phosphate group, the theoretical m/z of its mono-, di- or tri-derivatized precursor is calculated (ΔM = n × 135.043, n = 1-3). To ensure high-confidence identification, we implemented a stringent three-part verification strategy: a match is accepted only when the predicted precursor m/z shift, a minimum MS^2^ spectral similarity threshold, and the presence of the enforced diagnostic fragment ([C_6_H_5_N_2_O_2_]^-^, m/z = 137.035) are simultaneously satisfied. This triple-lock mechanism enables the robust identification of derivatized products that evade conventional spectral libraries while fully retaining the sensitivity gain introduced by derivatization. This integration of experimental derivatization with a computationally processed theoretical database overcomes the long-standing identification bottleneck in derivatization-based metabolomics and represents the central innovation of the method.

**Figure 1.**
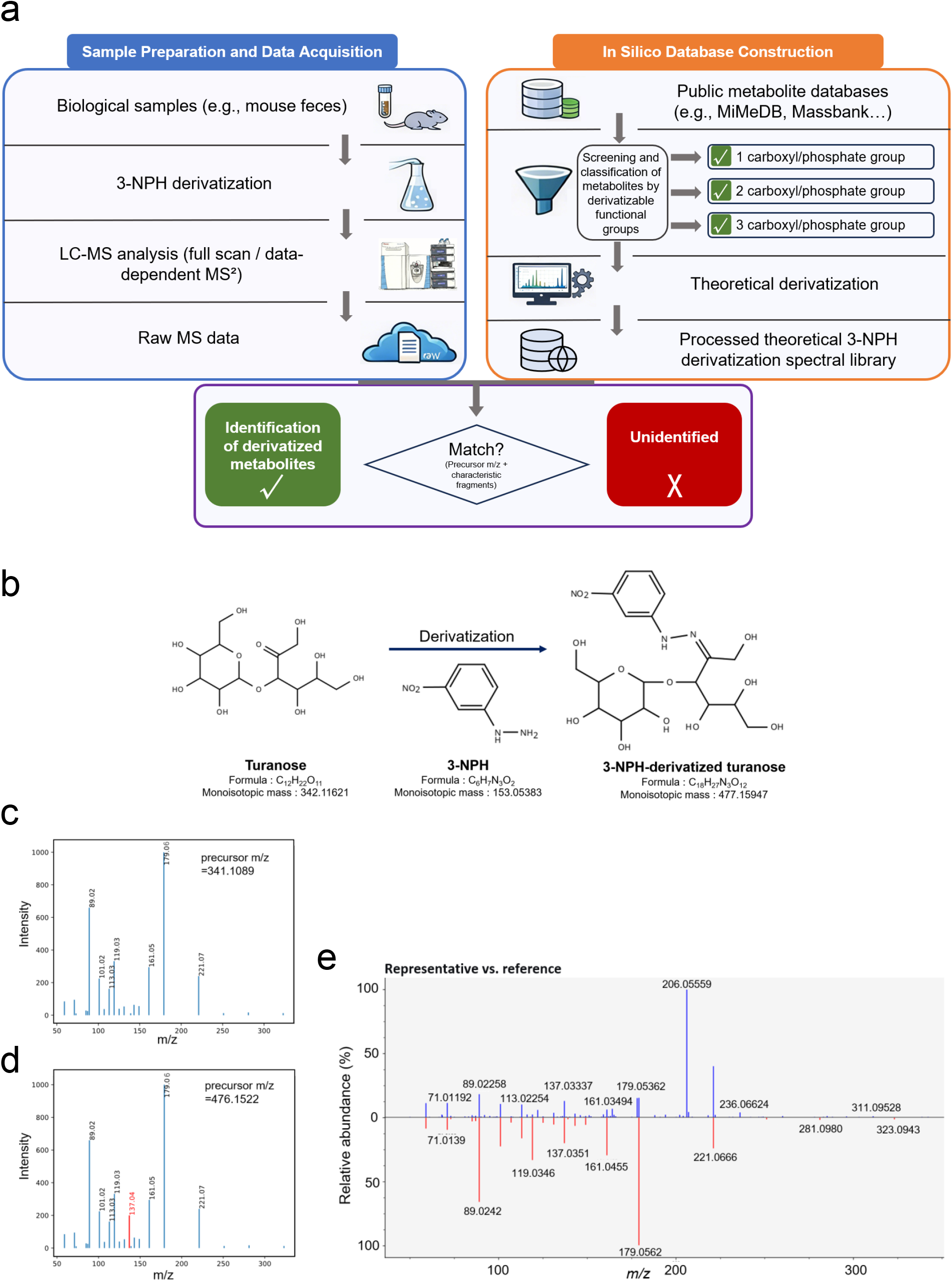
Establishment and analytical workflow of the 3-NPH derivatization-based untargeted metabolomics method. a, Overall workflow combining experimental derivatization with a computationally derivatized spectral library built from public databases by in silico mono-, di- and tri-derivatization; A match is accepted only when the predicted precursor m/z shift (ΔM = n × 135.043, n = 1-3), the MS^2^ spectral matching, and the enforced diagnostic fragment [C_6_H_5_N_2_O_2_]^-^ (m/z 137.035) are all satisfied. b, Example derivatization of turanose (C_12_H_22_O_11_, exact mass 342.1162): 3-NPH reacts with the reducing-end carbonyl to form a hydrazone, shifting the precursor by 135.043 and yielding derivatized turanose C_18_H_27_N_3_O_12_ (precursor m/z 476.1522). c,d, Mass spectra of native (precursor m/z 341.1089) and 3-NPH-derivatized turanose (precursor m/z 476.1522); derivatization introduces the characteristic diagnostic ion at m/z 137.035. e, Mirror-plot comparison of the experimental and in silico reference spectra, confirming identification.

To validate this theoretical library approach, we first examined turanose, a disaccharide containing a free carbonyl group that undergoes mono-derivatization (Fig. 1b). The original turanose spectrum in MassBank showed the characteristic fragmentation of the underivatized molecule (Fig. 1c). After in silico derivatization, we generated a predicted MS/MS spectrum featuring the diagnostic m/z 137.035 fragment alongside a mass-shifted precursor and product ions (Fig. 1d). When 3-NPH-derivatized turanose was analyzed by LC-MS/MS, the experimentally acquired spectrum (Fig. 1d, upper panel) showed near-perfect matching with the theoretically reconstructed spectrum (Fig. 1d, lower panel), including the m/z 137.035 diagnostic ion. This confirmed that our theoretical derivatization accurately predicts the MS/MS behavior of 3-NPH derivatives. Consistent with previous reports on 3-NPH derivatization, a side-by-side comparison of SCFA standards (Table 1, Supplementary fig.1) confirmed improved detection for low-molecular-weight analytes (e.g., FA 1:0-5:0) and a 51-to 232-fold response increase for FA 6:0-8:0, supporting the sensitivity advantage of this platform before its application to complex biological samples.

**Table 1.** Enhanced detection of short-chain fatty acids (SCFAs) by 3-NPH derivatization.

|  | Non-derivatized |  | 3-NPH-derivatized |  |
| --- | --- | --- | --- | --- |
|  | Peak profile | Response | Peak profile | Response |
| FA 1:0 | ND | - | Clear | 2.70E+08 |
| FA 2:0 | ND | - | Clear | 1.20E+08 |
| FA 3:0 | ND | - | Clear | 3.21E+06 |
| FA 4:0 | Noisy | - | Clear | 1.90E+07 |
| FA 5:0 | Noisy | - | Clear | 4.60E+06 |
| FA 6:0 | Clear | 1.58E+05 | Clear | 2.61E+07 |
| FA 7:0 | Clear | 8.74E+04 | Clear | 2.03E+07 |
| FA 8:0 | Clear | 2.45E+05 | Clear | 1.26E+07 |

*Short-chain fatty acid standards were prepared and analyzed at a uniform concentration of 5 $\mu$ g/mL. Data were extracted from the extracted ion chromatograms, and relative response represents the peak height of the target ion. ND, not detected. Noisy, no distinct peak discernible above background noise. Clear, well-defined peak.*

### Multi-compartment profiling reveals compartment-specific composition and restricted overlap of gut microbiota-related metabolites

Sixty samples (twelve mice × five compartments) were analyzed by the 3-NPH derivatization untargeted metabolomics platform, yielding 301 metabolite peaks (metabolite × derivatization state, including mono-, di- and tri-derivatized forms). Within each compartment, a metabolite was considered detected when its missing rate was below 70% across the 12 samples. Under this rule, 292, 294, 233, 255 and 248 metabolites were detected in the cecum contents, feces, serum, liver and brain, respectively (Fig. 2a-c). The gut compartments (cecum and feces) showed the broadest coverage (292-294), whereas serum showed the narrowest (233). Overall, the detected compounds spanned carboxyl-containing compounds, fatty acyls, organooxygen compounds, steroids, indoles, organic acids and bile acid structural classes, consistent with a derivatization reaction biased toward carboxyl, carbonyl and phosphate groups.

**Figure 2.**
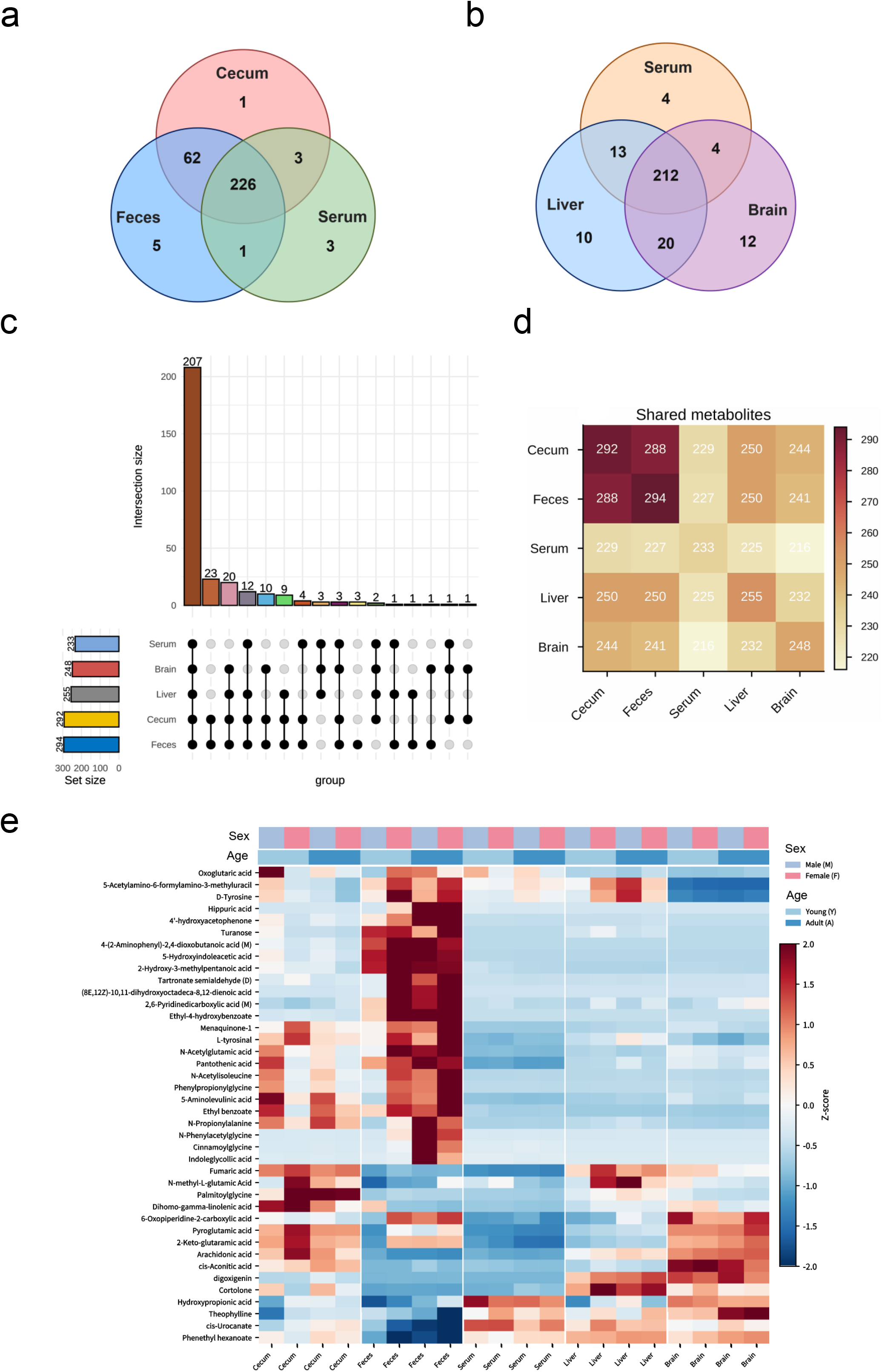
Compartment-specific composition and inter-compartment overlap of detected metabolites. a, Venn diagram of shared and compartment-specific metabolites among the gut-associated compartments (cecum contents, feces and serum). b, Venn diagram among serum, liver and brain. c, UpSet plot showing the overlap of detected metabolites across the five compartments. Left: total number of metabolites detected per compartment (set size). Top: number of metabolites in each intersection, with connected dots indicating the compartments contributing to that intersection. d, Shared metabolites between each pair of compartments. e, Heatmap of the most abundant metabolites across compartments and biological groups. Z-score-normalized mean peak area of the top 40 metabolites (rows, hierarchically clustered) across compartment, age (Y, 2-3 weeks; A, 8-10 weeks) and sex (M, male; F, female) subgroups (columns). Colour-coded annotation strips above and below the heatmap indicate the sex (M/F) and compartment of each column, and the age key; bars under the heatmap show compartment, age and sex identifiers for each sample column.

To assess how much of the metabolome is shared among compartments, we examined the overlap of detected metabolites. The cecum contents and feces showed the greatest overlap in detected features, consistent with their shared origin within the gut lumen, whereas the serum, liver and brain showed more restricted overlap with the gut compartments (Fig. 2d). Hierarchical clustering of the top-detected metabolites across compartments—z-score normalized across the compartment-by-age-by-sex groups for visualization only—grouped the two gut compartments together and separated them from peripheral compartments, while the brain and liver clustered more closely to one another (Fig. 2e). This compartment-resolved pattern indicates that although a shared core of microbiota-related metabolites reaches multiple organs, the peripheral metabolome is not a simple replica of the gut metabolome.

### Age- and sex-associated differences are compartment-specific

We compared individual metabolites between age groups and between sexes within each compartment (Mann-Whitney U test, unadjusted p < 0.05 and |log2 fold change| ≥ 1) (Fig. 3a-c). The number of age-associated metabolites was highest in feces (58) and serum (30), but lower in the cecum (10), liver (9) and brain (6). In contrast, sex-associated metabolites were highly localized, peaking in the gut (cecum: 54; feces: 54) but diminishing sharply in the serum (8), liver (1) and brain (1). These data indicate that sex-associated differences in the microbiota-related metabolome are most pronounced at the intestinal source and attenuate toward peripheral compartments, whereas age-associated differences are more broadly distributed. Notably, sex effects outnumbered age effects in the cecum (54 versus 10), while age effects dominated in the serum (30 versus 8).

**Figure 3.**
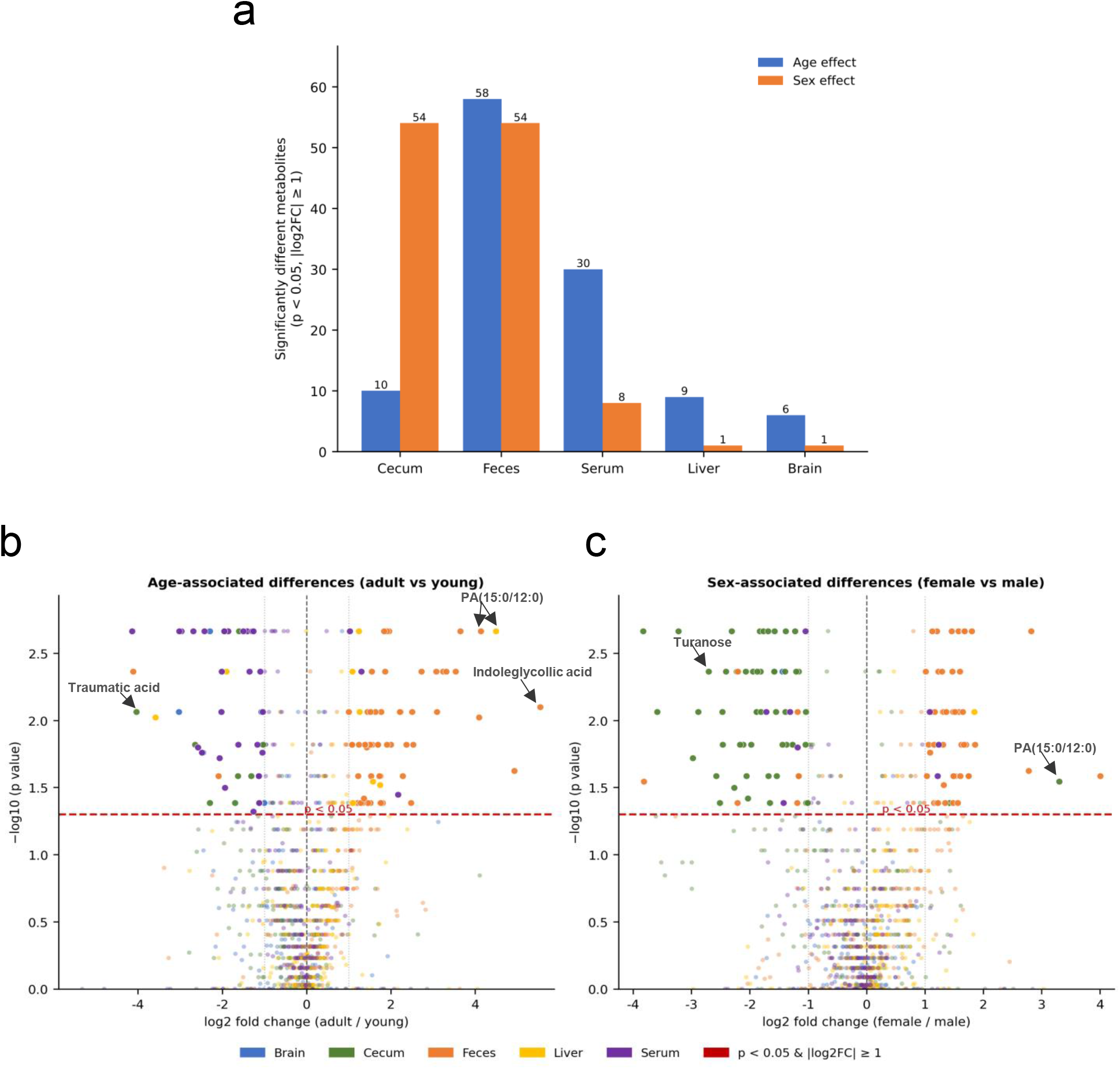
Compartment-specific age- and sex-associated differences in the microbiota-related metabolome. a, Number of significantly differing metabolites associated with age (blue) or sex (orange) within each compartment. Significance was determined by a two-sided Mann-Whitney U test (unadjusted p < 0.05 and |log_2_ fold change| ≥ 1). b,c, Volcano plots showing age- and sex-associated differences in metabolite abundance across compartments. Each point represents a metabolite-compartment pair with a missing rate < 50% and is coloured by compartment of origin. Significantly differing metabolites (two-sided Mann-Whitney U test, unadjusted p < 0.05 and |log_2_ fold change| ≥ 1) are depicted as larger points with higher opacity, whereas all other tested metabolites are shown as smaller, semi-transparent points. The red dashed line marks p = 0.05. b, Age-associated differences: log_2_ fold change (adult vs. young) on the horizontal axis against −log_10_(p value) on the vertical axis. c, Sex-associated differences: log_2_ fold change (female vs. male) against −log_10_(p value). Compartments are colour-coded: blue (brain), green (cecum), orange (feces), yellow (liver) and purple (serum). Data represent n = 6 mice per age × sex group.

This compartment-specific regulation was exemplified by several representative metabolites (Fig. 3b, c, Supplementary Table 1). Males exhibited prominent cecal enrichment of turanose [log2(female/male) = −2.58], whereas females showed higher cecal levels of PA(15:0/12:0) [log2(female/male) = 3.30]. Regarding age-dependent changes, adult mice had markedly higher fecal levels of indoleglycollic acid [log2(adult/young) = 5.54], while young mice showed elevated cecal traumatic acid [log2(adult/young) = −4.03]. Notably, PA(15:0/12:0) also increased with age in both feces [log2(adult/young) = 4.1] and liver [log2(adult/young) = 4.5], suggesting potential combined regulation by both age and sex.

### Coordinated metabolite pairs suggest candidate routes along the gut-brain and gut-liver axes

To explore potential coordination of gut microbiota-related metabolites across compartments, we computed pairwise Spearman correlations for each metabolite, aligning samples from the same mouse across compartments (12 matched mice) and imputing undetected (zero) values with half the minimum detected value of that metabolite before ranking. Because positive inter-compartment correlations are the most informative for coordinated changes in shared metabolites, we first selected features showing at least two significant (p < 0.05) positive cross-compartment correlations and visualized these features. For the selected metabolites, the complete Spearman correlation matrix across all ten compartment pairs (red, positive; blue, negative) is shown in Fig. 4a. Notably, several metabolites showed strong coordinated positive signals along the gut-brain and gut-liver axes, including the oligosaccharide D-Gal α1→6D-Gal α1→6D-glucose between cecum and brain (r = 0.76, p = 0.004), gamma-glutamylphenylalanine between cecum and brain (r = 0.69, p = 0.013), PA(15:0/12:0) between feces and liver (r = 0.83, p < 0.001), cinnamoylglycine between feces and liver (r = 0.79, p = 0.002) and phenethyl hexanoate between cecum and liver (r = 0.76, p = 0.005) (Fig. 4b-g). The positive pairwise correlations were additionally summarized for all ten compartment pairs in a chord diagram, in which each line represents one significantly positive cross-compartment correlation (p < 0.05, r > 0) for a single metabolite and its colour encodes that correlation’s Spearman coefficient r (Fig. 4h). Collectively, these coordinated positive gut-peripheral signals are consistent with shared origins and the potential transfer of gut microbiota-related metabolites along the gut-brain and gut-liver axes. In particular, the strong correlation of PA(15:0/12:0) implies a potential gut microbial contribution to the hepatic pool of odd-chain fatty acid (OCFA)-containing PA via the gut-liver axis.

**Figure 4.**
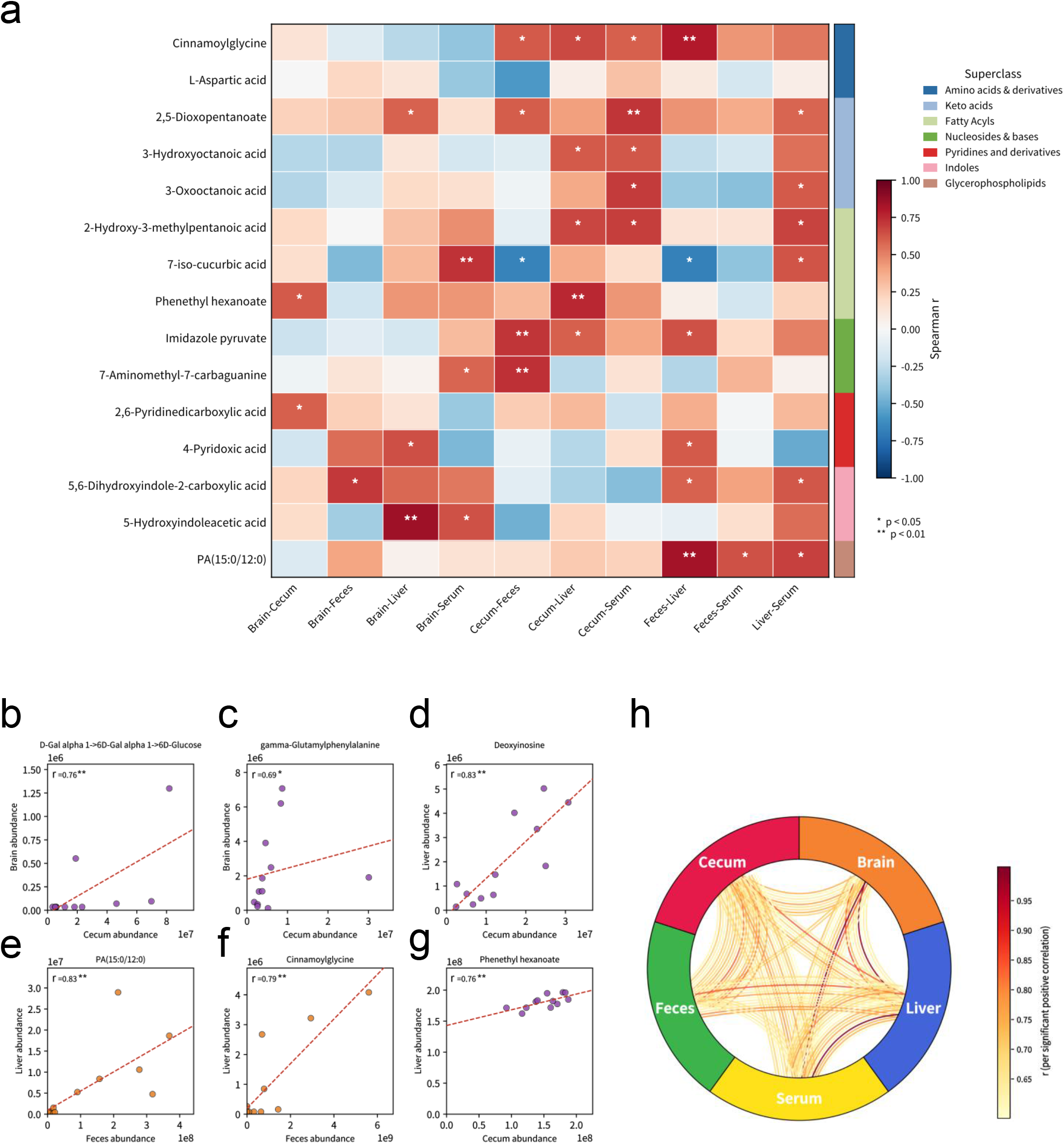
Cross-compartment correlation analysis along the gut-brain and gut-liver axes. a, Heatmap of the full Spearman correlation matrix (red, positive r; blue, negative r) for the 15 selected metabolites across all ten compartment pairs; metabolites are clustered by compound class on the vertical axis, and the right colour strip indicates the class of each metabolite. Asterisks mark the significance of each correlation (unadjusted p value; *, p < 0.05; **, p < 0.01). b-g, Representative gut-brain and gut-liver scatter plots with best-fit lines for selected literature- and research-highlighted metabolites, provided as six individual panels (b, D-Gal α1→6D-Gal α1→6D-glucose, cecum-brain; c, gamma-glutamylphenylalanine, cecum-brain; d, deoxyinosine, cecum-liver; e, PA(15:0/12:0), feces-liver; f, cinnamoylglycine, feces-liver; g, phenethyl hexanoate, cecum-liver). h, Chord diagram of the positive cross-compartment correlations for all ten compartment pairs; each line corresponds to one significantly positive correlation (unadjusted p < 0.05, r > 0) for an individual metabolite and is coloured by that correlation’s Spearman r.

## Discussion

In this study, we developed an untargeted metabolomics method based on 3-NPH derivatization coupled with a purpose-built theoretical spectral database. Applying this highly sensitive platform to five biological compartments (cecal contents, feces, serum, liver and brain) from young and adult mice of both sexes provided a compartment-resolved view of the gut microbiota-related metabolome. By integrating derivatization-driven sensitivity gains with the theoretical library, this approach detects a broad range of microbiota-related compounds— including low-molecular-weight organic acids that evade conventional untargeted analysis^7,8^.

Two conceptual findings emerge. First, gut-associated compartments (cecum, feces) carried the widest metabolome, whereas the circulation (serum) contained a smaller and partially overlapping subset, and the brain and liver harboured yet more restricted but compartment-specific pools. This is compatible with the view that while many gut-derived compounds enter the circulation, the peripheral metabolome is shaped by compartment-specific uptake, metabolism and barrier functions rather than being a passive mirror of the gut^6,9^. Second, sex-associated differences diminished markedly from the gut to the serum, liver and brain (54 and 54 vs. 8, 1 and 1). This gradient suggests that sex influences the microbial metabolome most strongly at its source in the gut, while peripheral filtering dampens sex effects downstream^10,11^. A recent multi-compartment metabolomics study similarly reported that while sex explained the least amount of variation at all body sites, its impact was significant in most compartments except the ileum, with the strongest sex effects observed in the gastrointestinal tract^12^. Mechanistically, sex hormones—particularly androgens—can directly shape gut microbial composition^13^, and the gut microbiota in turn regulates the enterohepatic circulation of androgens via β-glucuronidase-mediated deconjugation^14^, establishing a bidirectional sex-microbiota-host metabolic axis that is most active at the intestinal source. As metabolites transit from the gut lumen into the circulation and peripheral organs, tissue-specific uptake, hepatic first-pass metabolism and blood-brain barrier selectivity progressively attenuate the sex-associated signals, consistent with the diminishing sex effects we observed along the gut-serum-liver-brain gradient. Age effects, by contrast, were compartment-dependent^15,16^. In a comprehensive assessment of microbiome, age and sex contributions to the mouse metabolome, age was the primary driver of variation in the liver and spleen, whereas microbiota and age explained similar proportions of variation in urine, serum and peritoneal fluid^12^. This compartment-specific age effect likely reflects the cumulative impact of aging on organ-specific metabolic pathways—such as hepatic lipid and bile acid metabolism—that are superimposed upon, but distinct from, microbiota-driven changes.

Correlation along the gut-brain and gut-liver axes suggested candidate routes for metabolite transport^4,5^. Because positive inter-compartment correlations are the most informative for the coordinated transfer of shared metabolites, we focused on features with strong positive cecum-brain, cecum-liver and feces-liver correlations. The oligosaccharide D-Gal α1→6D-Gal α1→6D-glucose and gamma-glutamylphenylalanine showed strong positive cecum-brain correlations, whereas PA(15:0/12:0), cinnamoylglycine and phenethyl hexanoate were strongly positively correlated between gut compartments and liver, consistent with coordinated transfer along the gut-liver axis. In parallel, age- and sex-associated differences were largely compartment-specific, with the gut metabolome more strongly affected than the brain, suggesting that peripheral metabolic changes do not automatically propagate to the central nervous system^17^. Here, we emphasize that cross-compartment correlation is suggestive of potential inter-compartment transport; establishing causality requires further tracer or perturbation experiments.

Among these candidates, PA(15:0/12:0) is of particular interest because its 15:0 acyl chain corresponds to pentadecanoic acid (C15:0), a microbiota-associated odd-chain fatty acid (OCFA). C15:0 is produced by several Bacteroidota genera, including Bacteroides uniformis, and C15:0 supplementation reduced atherosclerotic plaque burden by approximately 50% in mice by inhibiting hepatic 3-hydroxy-3-methylglutaryl-CoA (HMG-CoA) reductase and enhancing low-density lipoprotein (LDL) clearance^18^. Absorbed long-chain fatty acids reach the liver in chylomicrons and albumin-bound pools, and microbe-derived odd-chain lipids are taken up by hepatosplanchnic tissues^19^. In hepatocytes, fatty acids are esterified into phosphatidic acid (PA), the Kennedy-pathway branch point for diacylglycerol and CDP-diacylglycerol synthesis^20^. However, our results suggest that hepatic esterification may not be the only plausible source. Commensal Bacteroides species possess the complete enzymatic machinery to synthesize PA de novo as a key intermediate of membrane phospholipid biosynthesis^21^. Moreover, spatially resolved lipidomics has shown that PA produced by Bacteroides thetaiotaomicron is detectable in the mouse colon, co-localizes with host tissues and shows the strongest host-transfer signal among the bacterial lipids examined^22^. Together, these findings support a direct bacterial origin for the PA(15:0/12:0) measured in our gut compartments, which contributes to the local PA pool and, upon absorption, to the circulating and hepatic PA pools.

A technical limitation of the current approach should also be noted. Particularly for small-molecule compounds that yield sparse MS/MS fragmentation in their native forms, derivatization can substantially alter the original fragmentation patterns. This alteration arises because the 3-NPH derivatization introduces a prominent moiety that shifts the primary fragmentation pathways away from cleavages driven by native derivatizable groups (e.g., carboxyl, carbonyl, and phosphoryl). Consequently, the derivatized structures predominantly generate reagent-specific diagnostic ions (most notably m/z 137, 152, and 122) that can dominate the product-ion spectra^23^. Additionally, the entire 3-NPH moiety may also be lost as a neutral fragment (153 Da)^8^, further remodeling the spectral landscape. This may result in lower spectral similarity scores (e.g., dot product) between the experimentally acquired spectra and the theoretical library entries, potentially increasing the risk of false negatives during library matching. To ensure effective and accurate identification of such compounds, we recommend two complementary strategies: first, whenever available, using derivatized authentic standards to build an empirical library for these specific analytes; second, when standards are unavailable, applying a more lenient spectral similarity cutoff (e.g., a dot product score of 0.3) while simultaneously imposing a more stringent requirement on the number of matched fragment ions (e.g., a minimum of three matched fragments). This combined scoring strategy can balance the inherent spectral alterations introduced by derivatization with the need for robust identification confidence. Moreover, given the limited applicability of in silico prediction tools (e.g., CFM-ID) for out-of-domain derivatized structures, generating full theoretical spectra entirely via computation was not adopted here^24^.

In conclusion, we developed a 3-NPH derivatization-based untargeted metabolomics platform with a purpose-built theoretical spectral library, overcoming the identification bottleneck in derivatization-based metabolomics. Applied across five mouse biological compartments, it resolved the gut microbiota-related metabolome, revealed that sex-associated differences were most pronounced in the gut and attenuated in peripheral compartments, uncovered compartment-specific age effects, and identified candidate transmission chains along the gut-brain and gut-liver axes, which warrant causal validation.

## Methods

### Animals

Twelve specific-pathogen-free (SPF) mice were purchased from Shanghai Xihe Biotechnology Co., Ltd. The animals comprised three males and three females at 2-3 weeks of age (young) and three males and three females at 8-10 weeks of age (adult). From each mouse, five biological compartments were collected: cecum contents, feces, serum, liver and brain, yielding 60 samples in total. All samples were stored at −80 °C until analysis.

### Reagents and chemicals

3-Nitrophenylhydrazine (3-NPH, ≥98% purity), 1-ethyl-3-(3-dimethylaminopropyl) carbodiimide (EDC, ≥98% purity), methanol (LC-MS grade), acetonitrile (LC-MS grade), pyridine (LC-MS grade) and ammonium acetate (LC-MS grade) were obtained from Sigma-Aldrich (St. Louis, MO, USA). Short-chain fatty acid (SCFA) reference standards (FA 1:0 to FA 8:0) were purchased from BePure (Beijing, China). Ultrapure water was generated using a Milli-Q system.

### Sample preparation and derivatization

Solid and tissue samples were processed identically. For cecal contents, feces, liver and brain, 10 mg of each sample was combined with two steel beads and 25 μL of ultrapure water, and homogenized at −10 °C for 3 min. Pre-chilled methanol (120 μL) was then added and homogenization was repeated at −10 °C for 3 min. After centrifugation at 18,000g for 20 min at 4 °C, 30 μL of the supernatant was transferred to a clean tube, with the remaining supernatant stored at −80 °C for further use. For derivatization, 20 μL of 3-NPH reagent solution (200 mM in methanol/water) and 20 μL of EDC solution (120 mM in methanol/pyridine) were added, and the mixture was incubated at 30 °C with shaking at 1,200 rpm for 1 h in a preheated metal bath. The reaction was quenched by addition of 130 μL of pre-chilled 50% methanol and centrifuged at 18,000g for 20 min at 4 °C (or at 4,000g for 30 min for 96-well plates). The supernatant was transferred to an autosampler vial for LC-MS analysis.

For serum samples, 25 μL of serum was mixed with 120 μL of methanol and vortexed. Metabolites were extracted with shaking at 1,200 rpm for 10 min at 4 °C, followed by incubation at −20 °C for 20 min and centrifugation at 18,000g for 20 min at 4 °C. Then 30 μL of the supernatant was transferred to a clean tube, with the remaining supernatant stored at −80 °C for further use, and derivatization was performed as described above (20 μL of 3-NPH reagent solution, 20 μL of EDC solution, 30 °C, 1,200 rpm, 1 h), followed by quenching with 130 μL of pre-chilled 50% methanol and centrifugation at 18,000g for 20 min at 4 °C. The supernatant was transferred to an autosampler vial for LC-MS analysis.

### LC-MS untargeted analysis

The analysis followed the protocol described in our granted invention patent^25^. Chromatography was performed on a Thermo Scientific UltiMate 3000 high-performance liquid chromatography (HPLC) system equipped with a Waters Acquity UPLC HSS T3 column (2.1 mm × 100 mm, 1.8 μm) at 40 °C, with a flow rate of 0.3 mL/min and an injection volume of 5 μL. Mobile phase A consisted of 10 mM ammonium acetate in water, and mobile phase B was acetonitrile. The gradient (v/v %B) was: 0-1 min, 1% B; 1-4 min, 1-48% B; 4-10 min, 48-100% B;

10-13.4 min, 100% B; 13.4-13.5 min, 100-1% B; 13.5-15 min, 1% B. Mass spectra were acquired on a Q Exactive Plus hybrid quadrupole-Orbitrap mass spectrometer in negative ESI mode (spray voltage, −4 kV; capillary temperature, 320 °C; sheath gas, 40 arb; auxiliary gas, 15 arb), with a scan range of m/z 170-2500 for derivatized samples, at a resolution of 70,000 (MS^1^) and 17,500 (MS^2^) in full-scan/data-dependent MS^2^ (Full MS/dd-MS^2^) mode.

Raw data were processed in MS-DIAL and searched against a purpose-built theoretical derivatized spectral database. The database was constructed by (i) retrieving structures of all metabolites from public negative-mode spectral libraries (MiMeDB^26^ and BILELIB^27^); (ii) selecting metabolites containing at least one 3-NPH-reactive functional group (carboxyl, carbonyl or phosphate), generating candidate lists for mono-, di- and tri-derivatization; (iii) computing the theoretical precursor shift upon reaction with n 3-NPH molecules, ΔM = n × (153.05383 − 18.01056) = n × 135.04327 (n = 1, 2, 3), giving precursor m/z [M − H + n × 135.04327]; and (iv) enforcing the characteristic 3-NPH diagnostic fragment [C□H□N□O□]− (m/z 137.035) in every theoretical MS/MS spectrum. For database searches, a match was accepted only when the MS/MS spectrum contained m/z 137.035 ± 0.02, with precursor and fragment mass tolerances of 0.01 Da.

Quantification was based on raw MS peak areas. For each compartment, a metabolite was considered detected if its missing values across the 12 samples was below 70%; otherwise it was treated as not detected in that compartment. For clustering and visualization only, missing values of detected metabolites were replaced by half the minimum detected value of that metabolite.

## Statistical analysis

Age- and sex-associated differences were assessed separately within each compartment. Both age and sex comparisons used two-group tests (Mann-Whitney U test) and reported unadjusted p values. For cross-compartment analysis, Spearman correlation was computed for each metabolite, using samples from the same mouse across the two compartments of each pair. All statistical analyses were performed using Python (NumPy, SciPy, pandas), R (version 4.5) within RStudio, and the Bioprofile Cloud platform.

## Data availability

The sample metadata and processed peak-area matrix are provided as Supplementary Table 2 and 3. The raw LC-MS/MS data are available from the corresponding author upon reasonable request.

