## Supplementary Fig. 1 for "Sensitive untargeted metabolomics via 3-nitrophenylhydrazine derivatization maps gut microbiota-related metabolites across multiple mouse biological compartments"

### Supplementary Figure

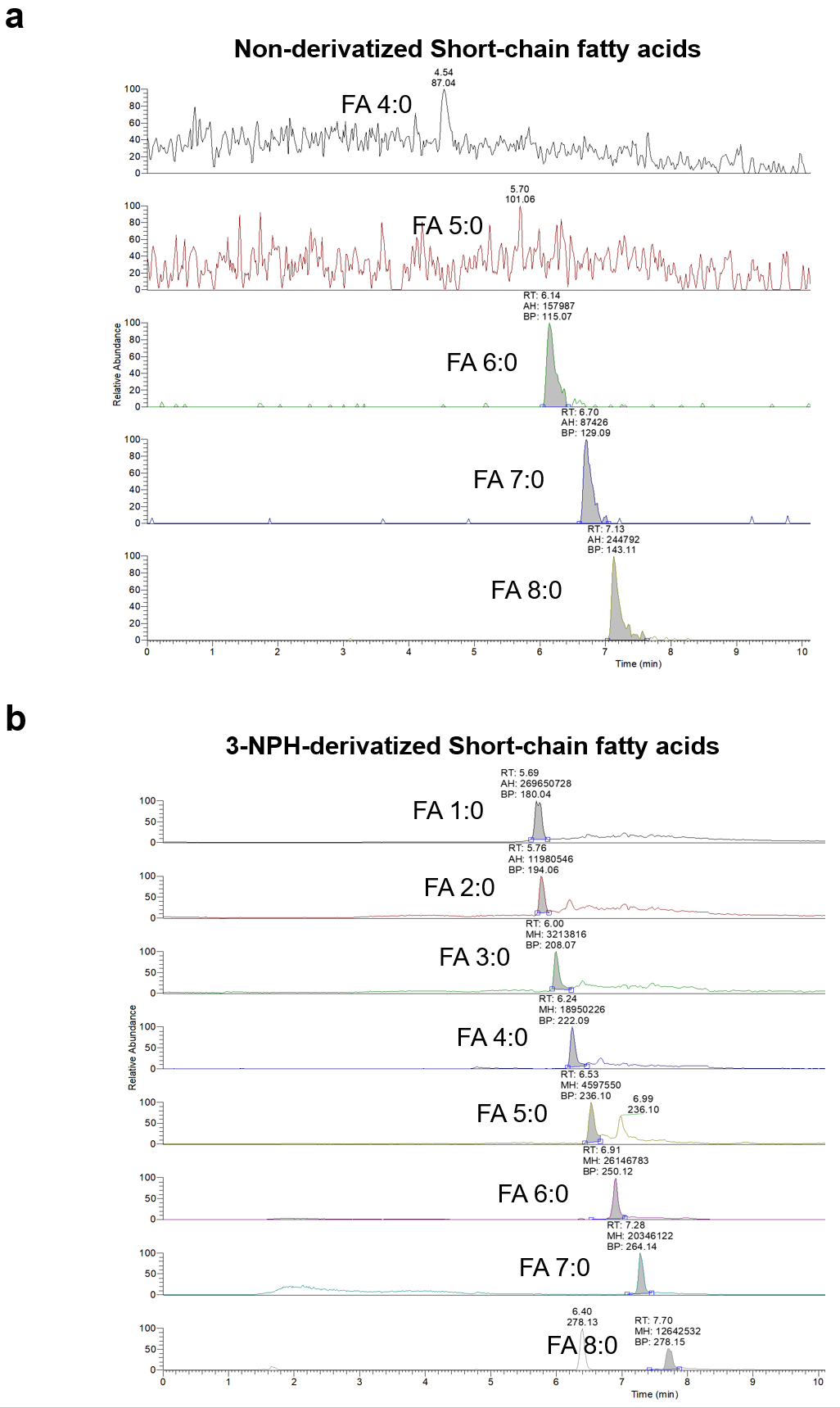

### Supplementary Fig. 1. Chromatographic comparison and sensitivity enhancement of short-chain fatty acid (SCFA) standards before and after 3-NPH derivatization. a. Extracted ion chromatograms (EICs) of non-derivatized SCFA standards (FA 4:0 to FA 8:0). Note the severe baseline noise and poor peak resolution for low-molecular-weight analytes (e.g., FA 4:0 and FA 5:0). b. EICs of 3-NPH-derivatized SCFA standards (FA 1:0 to FA 8:0). Derivatization enabled the stable detection of highly volatile, low-molecular-weight SCFAs (FA 1:0–3:0) and significantly improved the signal-to-noise ratios. Compared to the non-derivatized counterparts, the derivatized FA 6:0–8:0 exhibited a 51- to 232-fold increase in peak response, demonstrating the substantial sensitivity advantage of the 3-NPH platform. RT, retention time; BP, base peak *m/z*; AH, peak area.
